# The DNA Damage-Sensing NER Repair Factor XPC-RAD23B Does Not Recognize Bulky DNA Lesions with a Missing Nucleotide Opposite the Lesion

**DOI:** 10.1101/2020.09.14.293290

**Authors:** Katie M. Feher, Alexander Kolbanovskiy, Alexander Durandin, Yoonjung Shim, Jung-Hyun Min, Yuan Cho Lee, Vladimir Shafirovich, Hong Mu, Suse Broyde, Nicholas E. Geacintov

**Affiliations:** Chemistry Department and Department of Biology, New York University, 100 Washington East, New York, NY 10003-5180; Department of Chemistry & Biochemistry, Baylor University, Waco, TX 76706, USA

## Abstract

The Nucleotide Excision Repair (NER) mechanism removes a wide spectrum of structurally different lesions that critically depend on the binding of the DNA damage sensing NER factor XPC-RAD23B (XPC) to the lesions. The bulky mutagenic benzo[*a*]pyrene diol epoxide metabolite-derived *cis*- and *trans*-B[*a*]P-dG lesions (G*) adopt base-displaced intercalative (*cis*) or minor groove (*trans)* conformations in fully paired DNA duplexes with the canonical C opposite G* (G*:C duplexes). While XPC has a high affinity for binding to these DNA lesions in fully complementary double-stranded DNA, we show here that deleting only the C in the complementary strand opposite the lesion G* embedded in 50-mer duplexes, fully abrogates XPC binding. Accurate values of XPC dissociation constants (*K*_*D*_) were determined by employing an excess of unmodified DNA as a competitor; this approach eliminated the binding and accumulation of multiple XPC molecules to the same DNA duplexes, a phenomenon that prevented the accurate estimation of XPC binding affinities in previous studies. Surprisingly, a detailed comparison of XPC dissociation constants *K*_*D*_ of unmodified and lesion-containing G*:Del complexes, showed that the *K*_*D*_ values were ∼ 2.5 – 3.6 times greater in the case of G*:Del than in the unmodified G:Del and fully base-paired G:C duplexes. The origins of this unexpected XPC lesion avoidance effect is attributed to the intercalation of the bulky, planar B[*a*]P aromatic ring system between adjacent DNA bases that thermodynamically stabilize the G*:Del duplexes. The strong lesion-base stacking interactions associated with the absence of the partner base, prevent the DNA structural distortions needed for the binding of the BHD2 and BHD3 β–hairpins of XPC to the deletion duplexes, thus accounting for the loss of XPC binding and the known NER-resistance of G*:Del duplexes.

## 1. Introduction

Nucleotide Excision Repair (NER) is an important cellular defense mechanism that removes genotoxic DNA lesions from cellular DNA by a complex multi-step process that requires ∼30 proteins.[1] It has been known for some time that the efficiencies of repair of structurally different DNA lesions by NER mechanisms are highly variable[2] and that some DNA lesions are fully resistant to NER.[3] Using one-dimensionally stretched DNA molecules and single molecule imaging techniques, it was shown that the DNA lesion recognition factor XPC-RAD23B (XPC),[4] or its yeast homolog Rad4-Rad23,[5] search for DNA lesions by a constrained one-dimensional diffusion mechanism. The XPC/Rad4 DNA damage-sensing complex recognizes the distortions of the DNA duplex associated with the lesion, rather than the lesion itself and binds to the site of the adduct.[6, 7]The formation of the XPC-damaged DNA complex is followed by the recruitment of TFIIH,[1, 8] a factor that contains ten proteins, including the helicases XPD and XPB that verify the presence of an authentic DNA lesion.[8, 9] A slowing or stalling of the helicase XPD results in the signal that recruits other NER factors to the site of the lesion including the endonucleases that incise the damaged strand on the two sides of the lesions.[10] This work shows that a pair of stereochemically related bulky DNA lesions that are moderate to excellent NER substrates in fully base-paired DNA duplexes that are efficiently recognized by XPC, lose their XPC binding affinities and thus become NER-resistant[3] when the single nucleotide in the complementary strand opposite the lesion is deleted.

Among the best known and most widely studied DNA lesions are the UV radiation-induced *cis-syn* cyclobutane pyrimidine dimer (CPD) and 6-4 photoproducts (6-4 PP).[11, 12] The CPD lesion is first recognized by the DDB2-DDB1 heterodimer that recruits the XPC-RAD23B-Centrin 2 complex to the UV photolesions.[13, 14] However, neither DDB2-DDB1 nor Centrin 2 appear to be required for the recognition and excision of many bulky and helix-distorting adducts.[15, 16]

While XPC binding is a critical requirement for successful NER in the case of most bulky DNA lesions, there are relatively few direct comparisons of NER activities and XPC binding affinities. Correlations between NER and XPC binding affinities were established for DNA lesions such as CPD and 6-4 UV photoproducts,[7] C8-guanine adducts derived from aminofluorene (AF) and N-acetyl-aminofluorene (AAF),[17, 18] and covalent aristolochic acid-derived adenine adducts in double-stranded DNA.[19] We found earlier that the differences in NER efficiencies of various stereoisomeric benzo[*a*]pyrene-derived diol epoxide (B[*a*]PDE) – guanine lesions in double-stranded DNA (the (+)-*cis-* and (+)-*trans*-B[*a*]P-dG adducts G*, Fig. 1) were not correlated with XPC binding affinities.[20] However, in that work, the binding of multiple XPC molecules to the same DNA molecule, even at very low XPC concentrations, prevented the accurate determination of single-XPC molecule binding affinities. Here, we employed an undamaged DNA competitor and electrophoretic mobility shift assays to accurately determine XPC binding affinities of G*:Del duplexes, and the related fully base-paired G*:C duplexes for comparison. The G*:Del structures are formed during translesion synthesis catalyzed by Y-family polymerases that can give rise to frameshift mutations,[21-23] and are therefore biologically significant.

**Figure 1.**
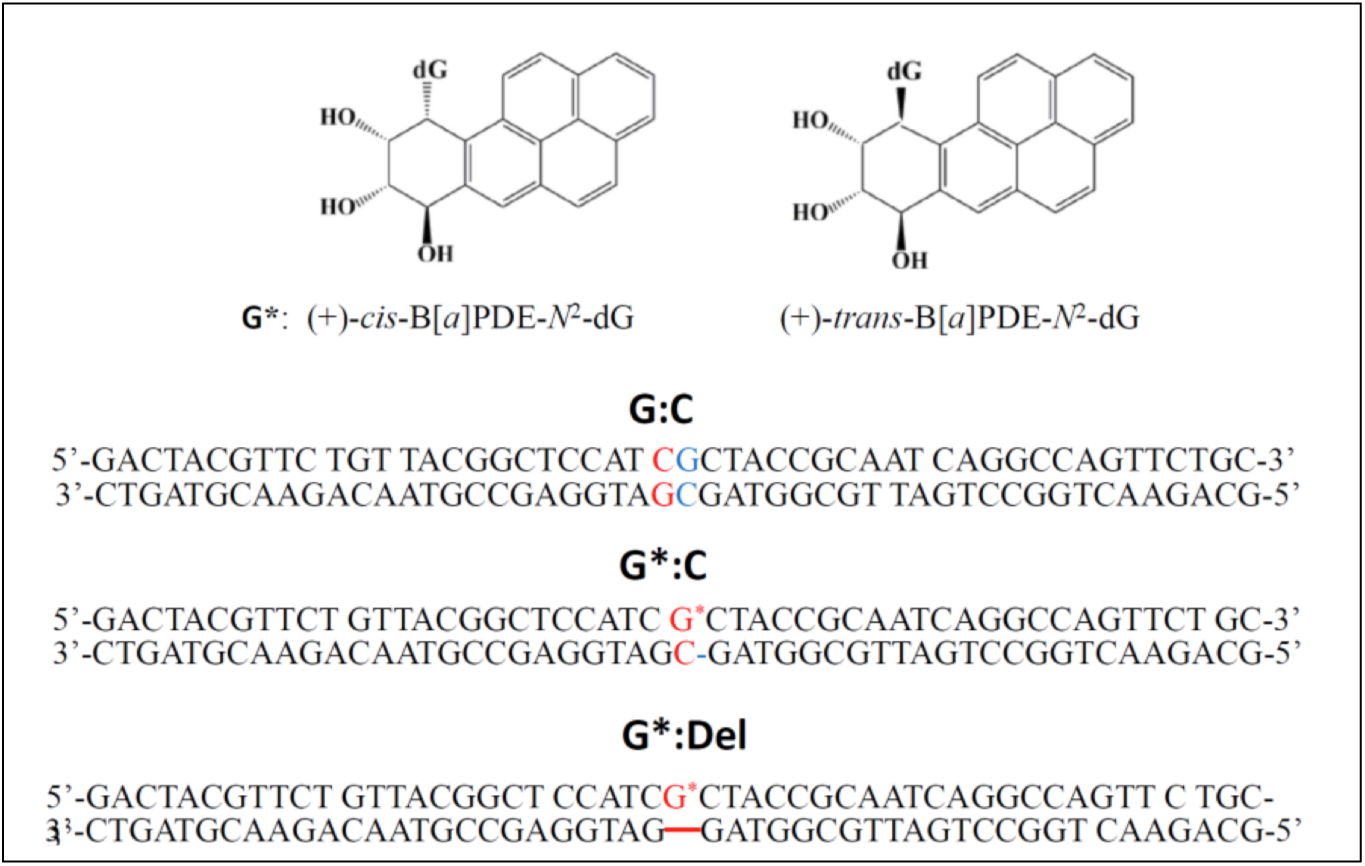
Stereochemical features of (+)-*trans*- and (+)-*cis*-B[*a*]P-dG (G*) DNA lesions, and sequences of unmodified G:C, full (G*:C) duplexes, and ‘deletion’ duplexes (G*:Del) containing single B[a]P-dG lesions.

The objectives of this work were to determine why DNA lesions that are moderate to excellent NER substrates, become fully NER resistant when embedded in G*:Del deletion duplexes.[24] Here we show that the deletion of the single complementary nucleotide C opposite (+)-*cis*- and (+)-*trans*-B[*a*]P-dG lesions embedded in 50-base pair DNA duplexes (Fig. 1), abrogates the specific binding of XPC to the G*:Del lesion (Table 1). This finding accounts for the NER resistance of the same lesions in G*:Del sequence contexts.

**Table 1.** Dissociation constants of single XPC-DNA complexes (*K*_*D*_), and of double (XPC_2_-DNA) complexes (*K*_*D2*_).

|  | Unmodified |  | *trans*-B[a]P-dG |  | *cis*-B[a]P-dG |  |
| --- | --- | --- | --- | --- | --- | --- |
|  | Full | Del | Full | Del | Full | Del |
| $K_D$, nM | $1.1 \pm 0.2$ | $1.2 \pm 0.2$ | $0.24 \pm 0.07$ | $2.6 \pm 0.30$ | $0.41 \pm 0.10$ | $3.6 \pm 0.60$ |
| $K_{D2}$, nM | $3.2 \pm 0.7$ | $3.6 \pm 0.7$ | --- | --- | --- | --- |

A rigorous analysis of the XPC concentration-dependent binding curves demonstrates that the dissociation constants *K*_*D*_ are 2.5 - 3.5 times greater than the *K*_*D*_ values of the unmodified G:C or G:Del duplexes defined in Fig. 1. These unusual and apparent DNA lesion avoidance effects are discussed and interpreted in terms of the structural features of Rad4/yeast XPC - G*:C complexes [7, 25] and Molecular Dynamics (MD) simulations. Finally, an intriguing observation is that a single G*:Del site positioned in the middle of a 50-mer duplex can strongly diminish the overall XPC binding affinity relative to the same lesion embedded in fully base paired G*:C and even unmodified G:C and G:Del duplexes.

## 2. Materials and Methods

### Preparation of modified 50-mer oligonucleotide duplexes and XPC proteins

The methods for generating these duplexes were described in detail earlier.[20] Briefly, 50-mer double-stranded sequences containing either the normal C opposite the modified guanine to form the unmodified G:C duplexes, or (+)-*trans*-or (+)-*cis*-B[*a*]P-dG (G*:C duplexes), or lacking the C nucleotide opposite the adduct in (G*:Del duplexes, Fig. 1), were prepared as described previously.[20] The B[*a*]P residue is linked to the exocyclic amino group of guanine positioned in the middle of in 50-mer the DNA sequences as shown in Fig. 1. The two 50-mer DNA duplexes were prepared by heating stoichiometric solutions of the two strands to 90 ^0^C, followed by slow overnight cooling of the solutions at 4 ^0^C. The duplexes were pre-purified by 12% polyacrylamide native gel electrophoresis. The full length XPC-RAD23B was expressed and prepared from insect cells using baculovirus overexpression system and chromatography methods similar to those described earlier.[6]

### Electrophoretic Mobility Shift Assays

The XPC binding studies were conducted in solutions containing 0.40 nM ^32^P-end-labeled 50-mer duplexes containing the B[*a*]P-dG (G*:C or G*:Del duplexes, Fig. 1) and 10 nM unmodified 50-mer duplexes that were used as competitor DNA. Employing this approach, the formation of multiple XPC molecules bound to the same DNA molecule[20] was minimized. Appropriate models of binding equilibria were utilized to fit the experimental XPC binding data, and to elucidate the dissociation constants *K*_*D*_.

The binding reactions were carried out in 8 µL aliquots of 0.40 nM ^32^P-end labeled DNA duplexes in 5 mM bis-tris propane-HCl, 75 mM sodium chloride, 10 mM dithiothreitol, 5% glycerol, 0.74 mM 3-[(3-cholamidopropyl)dimethylammono]-1-propanesulfonate (CHAPS), and 500 µg/mL bovine serum albumin (pH 6.8) buffer containing different XPC concentrations. Before each experiment, the activity of the XPC samples was verified. All samples containing 0.4 nM 32P-labeled 50-mer DNA duplexes and the same, but unmodified competitor DNA (10 nM), and XPC were incubated for 20 min at 23 ^0^C. Finally, the samples were loaded and electrophoresed on a 96.5 cm wide and 127 cm long 4.8% native polyacrylamide gel (acrylamide/bisacrylamide 37.5:1) at 500 V and 4° C for 50 minutes in a phosphate running buffer (25.5 mM H_2_PO4 and 24.5 mM KHPO4, pH 6.8). The gels were dried and scanned using a Typhoon FLA 9000 phosphorimager (GE Instruments). The relative fractions of bound and unbound DNA were evaluated using ImageQuant 5.2 software. All experiments were conducted at least three times.

## 3. Results

### 3.1. Background

It is well established that XPC is capable of binding to unmodified DNA duplexes[25, 26]. After specific complex formation with XPC, the helicase-driven verification mechanism involving TFIIH distinguishes between DNA with or without modifications or lesions, thus avoiding futile DNA excision events in the absence of a DNA lesion (e.g., in the case of mismatched DNA, or single-strand/double-strand junctions that are specifically recognized by XPC, but do not elicit NER). However, some DNA lesions are resistant to NER[3] which can occur either because of a failure of productive XPC binding, or a failure of the verification mechanism even when XPC binding to DNA lesions does occur. We have previously shown that XPC binds with high affinity to benzo[*a*]pyrene-derived B[*a*]P-dG*:C lesions embedded in full 50-mer duplexes with a canonical C opposite G*.[20] However, we show here for the first time that XPC surprisingly binds with significantly lower efficiencies to the same DNA lesions in G*:Del duplexes, displaying an even lower binding affinity than manifested by the lesion-free G:C and G:Del duplexes (Figs. 2 and 3).

**Figure 2.**
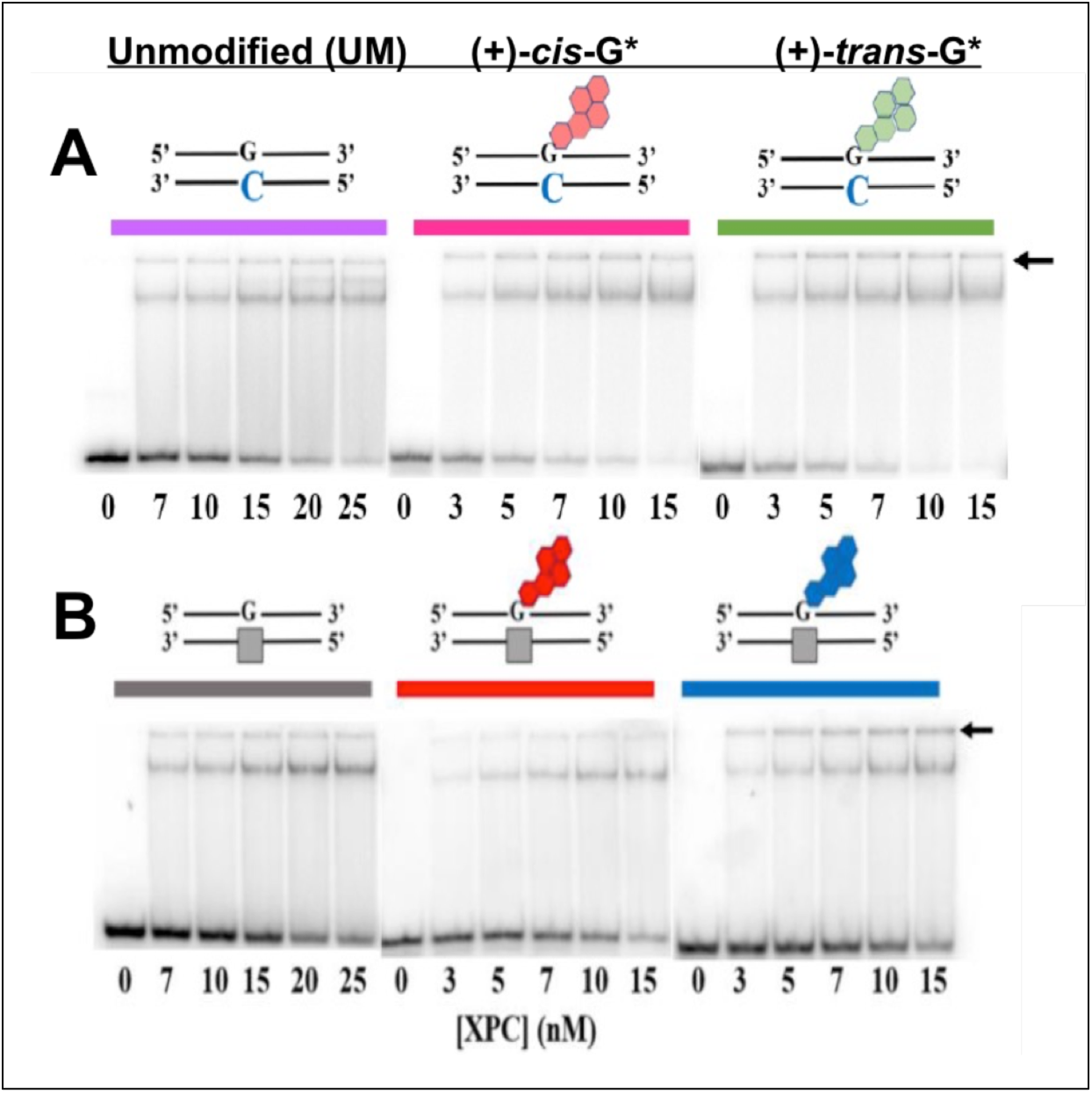
Gel electrophoresis mobility shift assays of XPC binding to unmodified (G:C) and B[*a*]P-modified (A) full (G*:C) 50-mer duplexes, and (B) deletion duplexes (G*:Del) defined in Figure 1.

**Figure 3.**
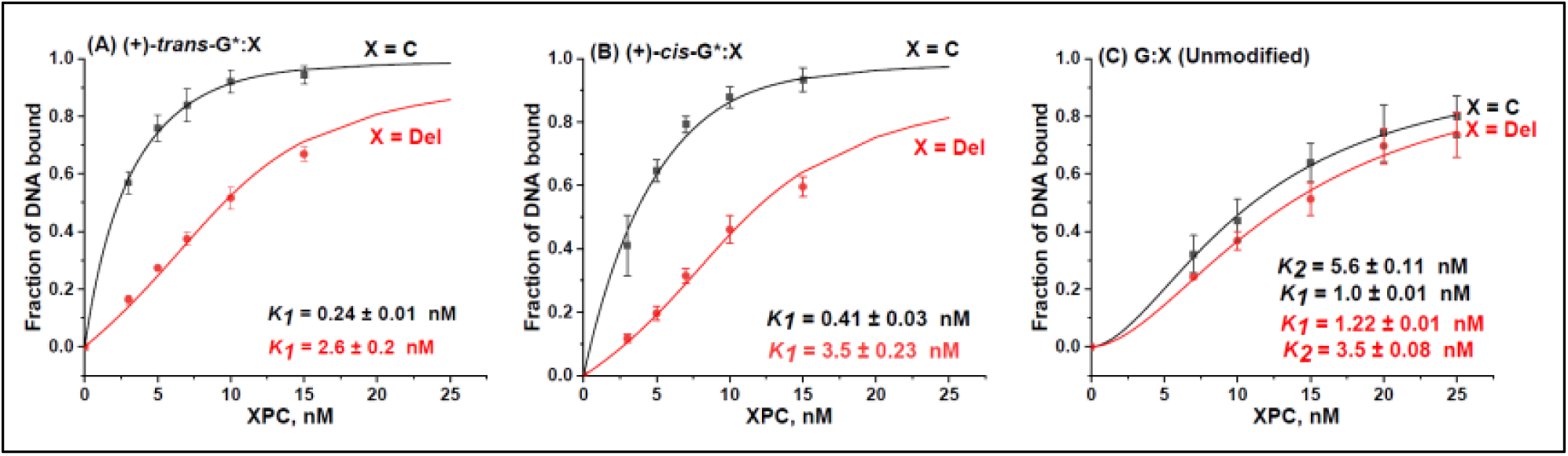
XPC binding to DNA duplexes with single ((A) (+)-*trans*-B[*a*]P-dG, (B) *cis-*B[*a*]P-dG (G*:C), and (C) unmodified full double-stranded DNA (G:C),) or deletion DNA duplexes (G*:Del or G:Del). The solid lines were computed as described in the text.

### 3.2. XPC–DNA complex formation

Typical EMSA results of XPC binding to unmodified 50-mer DNA duplexes G:C and G:Del, and to the same duplexes containing single B[*a*]P-dG lesions are depicted in Fig. 2, and the corresponding densitometry tracings are shown in Fig. 3. The stronger upper bands in the autoradiographs (Fig. 2) are due to the slower migrating XPC-DNA complexes, while the lower bands are due to XPC-free DNA duplexes. We have previously described the nature of the binding of XPC to full G*:C duplexes and found that XPC could form multimeric complexes with DNA [20] that made it difficult to accurately assess the equilibrium dissociation constants for 1:1 XPC:DNA binding. In the presence of a 25-fold higher concentration of unmodified G:C duplexes, single XPC-DNA bands were dominant in all experiments as illustrated in Fig.2. Minor amounts of the XPC-DNA complexes are trapped in the wells (the positions of the wells are denoted by the arrows in Fig. 2). The fractions of ^32^P-labeled XPC-DNA complexes as a function of XPC concentration are plotted in Figures 3A-C. A significant enhancement in XPC binding is observed in the B[*a*]P-modified full G*:C duplexes, consistent with our previous results that were conducted without unmodified competitor DNA. [20] In the case of unmodified DNA [G:C] and [G:Del] duplexes, the shapes of the binding curves are similar, and almost indistinguishable (Fig. 3C). This illustrates that a single unpaired dG that is missing its Watson-Crick partner in G:Del duplexes, does not provide sufficient distortion to 50-mer double-stranded DNA to be specifically recognized by XPC. Furthermore, XPC does not recognize either stereoisomeric B[*a*]P-dG lesion opposite the deletion site in [G*:Del] duplexes, thus demonstrating that the absence of the deoxycytidine residue[27] causes a remarkable impediment to XPC binding to the G*:Del site (Figs. 3A and 3B).

### 3.3. Analysis of XPC Binding to B[*a*]P-modified full and deletion duplexes

**3.3.1**. The following binding equilibria were employed in order to calculate the dissociation constants *K*_*D*_ = *k*_*d*_ */k*_*a*_, where *k*_*d*_ and *k*_*a*_ are the rate constants of dissociation and association of XPC-DNA complexes, respectively. The XPC binding to G*:C duplexes was analyzed in terms of the following standard model:

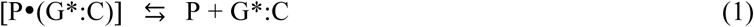

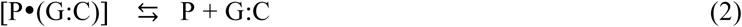

With dissociation constants

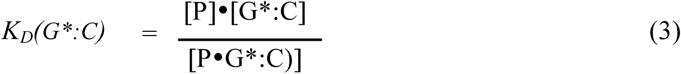

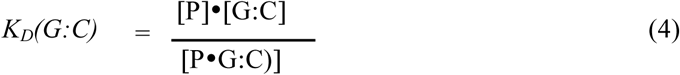

where [P] denotes the XPC concentration, and G:C and G*:C denote the 50-mer duplexes defined in Figure 1. In the case of G*:Del duplexes, the values for G*:C in equations (1)-(4) were replaced by G*:Del. The exact solutions of such coupled binding equilibria have been considered by Wang.[28] and were used here to evaluate the XPC dissociation constants *K*_*D*_(G*:C) and *K*_*D*_(G*:Del). The unmodified DNA dissociation constants *K*_*D*_(G:C) or *K*_*D*_(G:Del) were determined as described in section 3.3.2 below. The best fits to the experimental XPC binding data are shown in Figures 3A and 3B, and the *K*_*D*_ values are summarized in Table1.

**3.3.2**. In the case of XPC binding to unmodified G:C or G:Del duplexes (Figure 3C), there is no competitor DNA. Thus, the calculation of the dissociation constants *K*_*D*_(G:C) in eq. 4 were based solely on the equilibrium eq. 2, while that of *K*_*D*_(G:Del) was based on G:Del substituting for G:C in eqs. (2) and (4).

The fractions of XPC-bound DNA molecules are, in principle, obtained by solving a simple quadratic equation that yields the dissociation constants *K*_*D*_.[29] However, this approach does not provide a satisfying fit to the experimental binding curves (not shown). Instead, these observations suggest that a second XPC molecule can bind to an XPC-unmodified DNA complex to form (XPC)_2_-DNA complexes. Indeed, a second, weaker upper band is observable in the case of the unmodified G:C duplexes, just above the main upper XPC-(G:C) complexes at 20 nM and 25 nM XPC concentrations, (Fig. 3A). The appearance of these upper bands is accompanied by an accumulation of radioactive material between the upper and the lower bands in this panel. This is evident from the densitometry traces of the 15 nM XPC experiment (Fig. 4(A)), and becomes significantly more pronounced as the XPC concentration is increased to 25 nM (Supporting Information). This effect is not observable, to any significant extent, either in the case of the G:Del, G*:C or G*:Del duplexes (Figs. 3B,C,D). The appearance of radioactive material between the upper and lower bands is primarily attributed to the dissociation of (XPC)_2_-DNA complexes (without the B[*a*]P-dG lesions) during the electrophoresis experiments.

**Figure 4.**
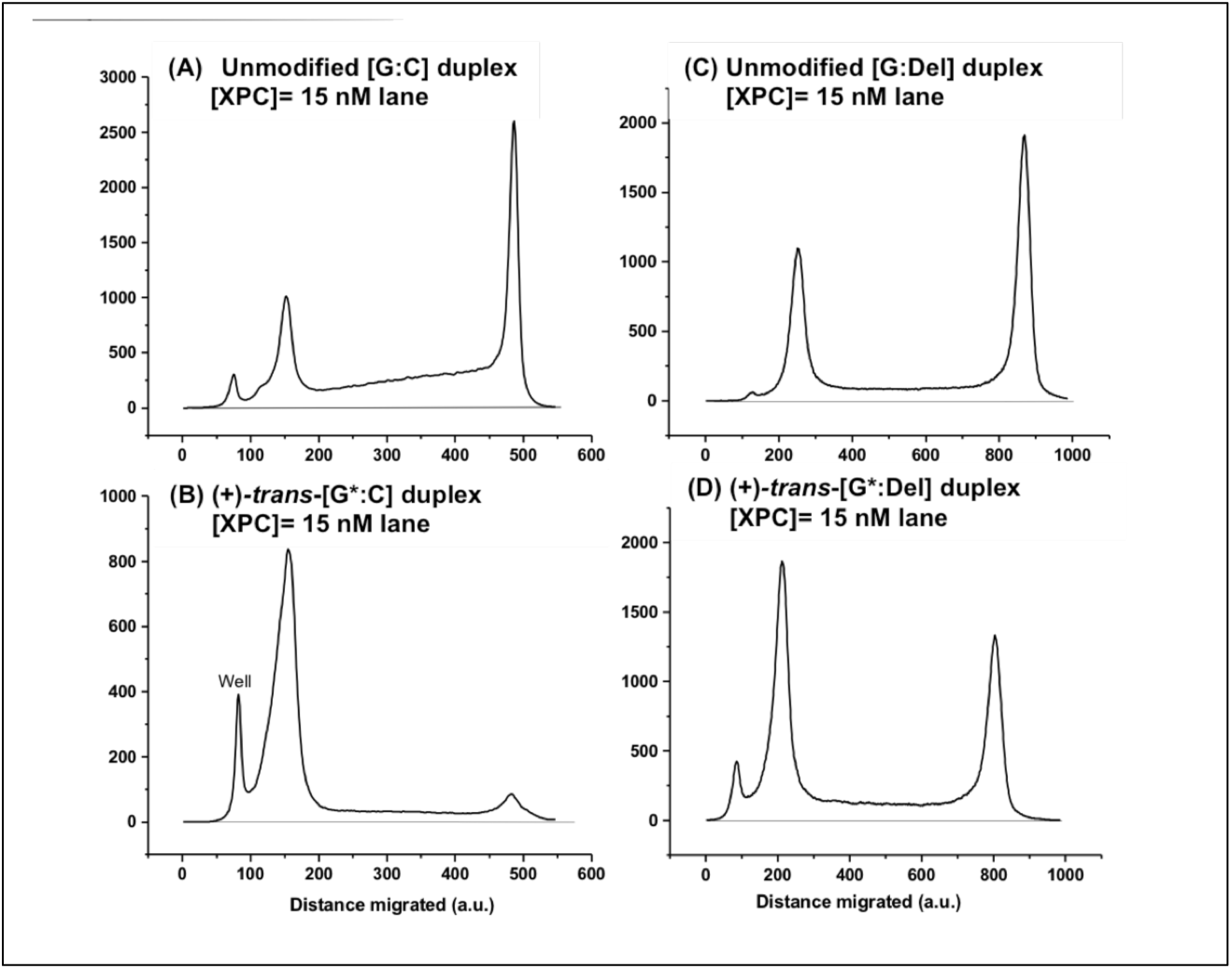
Densitometry tracings of selected (15 min) lanes shown in Figure 2.

The dissociation constants for the unmodified XPC-(G:C) and XPC-(G:Del) duplexes were evaluated by considering the equilibrium in Eq. 2, coupled with the following equilibrium and eq. 5:

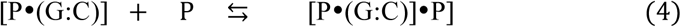

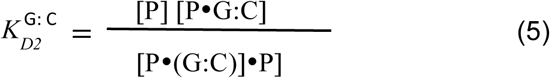

In order to compare the values of *K*_*D*_(G:C) and *K*_*D*_(G:Del), equations (3) and (5) were combined and the resulting cubic equations were solved to determine the overall fractions of DNA molecules associated with one or two XPC molecules using appropriate mathematical approaches (Supporting Information). The best values of these two dissociation constants that provide a good fit of the calculated XPC binding curves to the experimental data (Figures 2 & 3) are shown in Table 1.

## 4. Discussion

### 4.1 The occurrence of deletion mutations by translesion synthesis *in vitro* and in cellular environments

When high-fidelity DNA polymerases encounter DNA lesions, their progress is stalled, and they are replaced by one or more low-fidelity translesion synthesis (TLS) polymerases.[30, 31] Translesion synthesis catalyzed by Y-family polymerases can result in the incorrect pairing of the modified guanine (G*) with a mismatched nucleotide thus causing single base substitution mutations. Furthermore, strand slippage can form misaligned frameshift intermediates which result in a deletion mutation since the modified template base is skipped by the polymerase.[32] It has been shown that TLS occurs with B[*a*]P-dG adducts in DNA *in vitro*[33] and in eukaryotic cells.[34-36] The G*:Del duplexes are fully resistant to NER in cell-free DNA repair assays, although the same adducts embedded in full G*:C duplexes are moderate-to-excellent NER substrates in human cell extracts.[24, 37, 38]The remarkable NER resistance of B[*a*]P-dG adducts in G*:Del deletion sequence contexts was not fully understood. Here we demonstrate that the DNA damage sensor XPC binds to G*:Del sequences characterized by dissociation constant that are *greater* than the *K*_*D*_ values associated with unmodified G:Del and G:C duplexes (Table 1).

### 4.2 XPC-DNA dissociation constants

Previously, we employed a bimodal Hill equation to derive the apparent dissociation constants *K*_*D*_ ≈ 0.7±0.2 nM for both (+)-*trans*- and (+)-*cis*-B[*a*]P-dG*:C duplexes, and 1.2 ±0.7 nM for unmodified G:C 50-mer duplexes.[20] In the present study we utilized an improved binding assay employing unmodified competitor DNA to minimize the formation of higher-order complexes., The rigorous Wang approach [28] was adopted for calculating the dissociation constants *K*_*D*_(G*:C) in the presence of competitor DNA. More precise relative values of *K*_*D*_(G*:C) = 0.24±0.05 (*trans*) and 0.41±0.07 nM (*cis*) were thus obtained (Table 1). In the case of unmodified duplexes, the low XPC concentration value of *K*_*D*_(G:C) = 1.0±0.2 nM is attributed to the dissociation constant for the binding of the first XPC molecule, and *K*_*D2*_(G:C) = 5.6±0.8 for the second. Based on these values, it is concluded that the XPC binding affinities for the B[*a*]P-dG adducts embedded in 50-mer G*:C duplexes is 2 – 4 times greater than to the same, but unmodified G:C duplex, and ∼ 9 – 10 times greater than to the G*:Del duplexes.

### 4.3 Comparisons of XPC binding affities and conformations of B[*a*]P-modified G*:Del and G*:C duplexes

#### Structural features of the DNA lesions

Using high resolution NMR methods it was demonstrated that <u>in full</u> *trans-*B[*a*]P-G*:C duplexes, the B[*a*]P aromatic ring system is positioned in the minor groove,[39] while in the case of the *cis*-B[*a*]P-dG*-C duplexes, it assumes a base-displaced intercalative conformation.[40] Both duplexes are thermodynamically destabilized by the adducts.[3] The NMR structures of the full (+)-*trans*-G*:C,[39] (+)-*trans*-G*:Del,[41] full *cis-*G*:C[40] and *cis* G*:Del[42] duplexes are compared and described in Figure 5. An important point is the wedge shaped intercalation pocket in the (+) *cis* and (+) *trans* G*:Del duplexes that is due to the deletion of the cytosine residue opposite the lesion. The strong Van der Waals B[*a*]P aromatic ring system– base stacking interactions enhance the thermodynamic stabilities and melting points of the G*:Del duplexes and they are consequently DNA repair-resistant.[38, 43]

**Figure 5.**
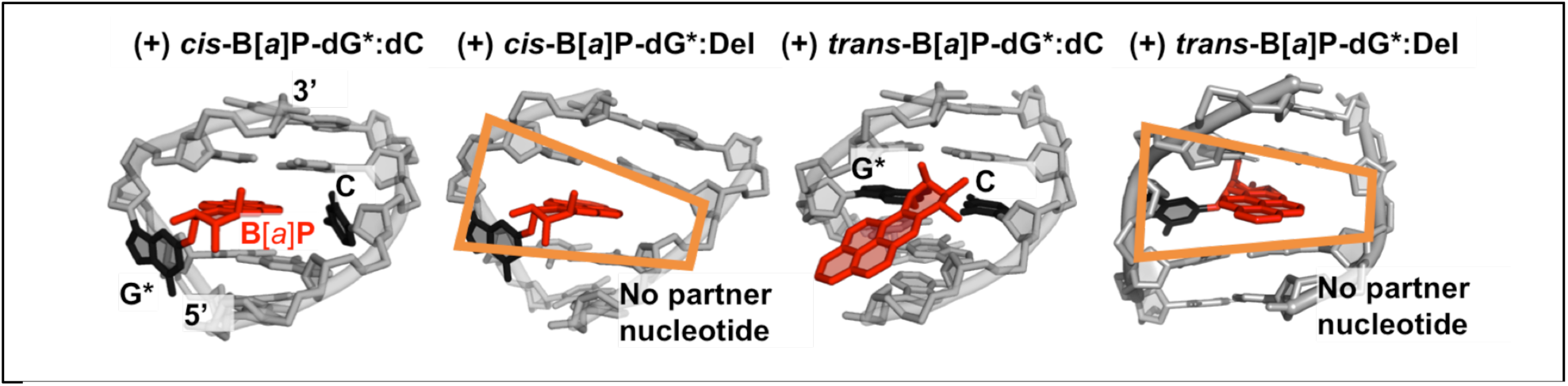
NMR solution structures for the (+) *cis*-B[*a*]P-G*:C (+) *cis*-B[*a*]P-G*:Del,(+) *trans*-B[*a*]P-G*:C and (+) *trans*-B[*a*]P-G*:Del duplexes. The (+) *cis*-B[*a*]P-G*:C is base-displaced intercalated with B[*a*]P rings intercalated into the duplex, the deoxyguanosine G* is displaced into the minor groove, and the partner C is displaced into the major groove. In the case of the (+) *cis*-B[*a*]P-G*:Del, the G* is displaced into the minor groove, and the absence of the partner C nucleotide gives rise to a wedge shaped intercalation pocket (orange frame) that enhances B[*a*]P aromatic ring stacking with adjacent base pairs. The full duplex (+) *trans*-B[*a*]P-G*:C has the aromatic ring system in the minor groove directed toward the 5’ end of the modified strand. The (+)-*trans*-G*:Del structure resembles the intercalated (+)-*cis*-G*:Del structure, except that the modified deoxyguanosine is displaced into the major groove (*trans*) whereas it is displaced into the minor groove in the case of the (+)-*cis* G*:Del lesion.

#### Dissociation constants

The higher dissociation constants of the two 50-mer B[*a*]P-G*:Del duplexes *K*_*D*_(G*:Del) ≈ 2.6 – 3.5 nM), relative to the analogous unmodified DNA G*:C duplexes (*K*_*D*_(G:C) ∼ 1.0 nM), indicates that XPC binds less strongly to the B[*a*]P-dG*:Del duplexes than to unmodified DNA by a factor of ∼ 2.5 - 3.5. It has been shown earlier that XPC does not recognize the UV photoproduct CPD, although the duplexes are thermodynamically destabilized.[12, 44][[25] By contrast, the G*:Del duplexes described here offer a new paradigm of lesion avoidance[45] by a bulky DNA lesion that stabilizes the local thermodynamic stability of the deletion site and therefore hinders strand separation.[38] Thus, the XPC protein, as it diffuses along the DNA duplex, is unable to attach to the site of the lesion in the deletion sequence context. It is striking that DNA lesion avoidance that enhances the XPC dissociation constant can occur when the partner nucleotide opposite the lesion is deleted in an otherwise fully base-paired DNA duplex 50 base pairs in length.

#### Unusual XPC lesion avoidance

The lesion avoidance mechanism may be understood at a molecular level in terms of the crystal structure of Rad4-Rad23B, the yeast ortholog of XPC-RAD23B that recognizes the distortions and destabilizations in double-stranded DNA caused by a DNA lesion[6, 7]. In these structures, the BHD2 β-hairpin of Rad4 binds to the minor groove, and the BHD3 β-hairpin inserts into the DNA to stabilize the Rad4/yeast XPC-DNA productive open complex, which allows the recruitment of subsequent NER factors. This structure, and the significant amino acid sequence and functional similarities between Rad4 and XPC, suggest that the β-hairpin insertion and local DNA strand separation are crucial elements for the recognition of DNA lesions by XPC as well.[6, 46]

#### MD simulations

The formation of the productive open complex occurs in stages that have been delineated in MD simulations: the BHD2 hairpin first binds to the DNA minor groove at the lesion site causing duplex unwinding and bending (initial binding); the partner base/bases then flip into the protein, and the BHD3 hairpin then inserts into the duplex.[46, 47] This was delineated for the case of the *cis*-B[*a*]P-dG*-C duplex that is efficiently excised by the NER system.[47] Molecular dynamics simulations of Rad4/yeast XPC and initial binding to lesion-containing DNA have explained structurally why the *cis*-B[*a*]P-dG*-Del duplex fails to bind with Rad4/yeast XPC. The NER resistant *cis*-B[*a*]P-dG*-Del duplex does not manifest any of the structural and dynamic features for successful initial binding: it remains essentially undisturbed by Rad4/yeast XPC without BHD2 hairpin minor groove binding, unwinding, bending, and partner base flipping that are needed to ultimately achieve the productive open complex[47] (Figure 6) A likely underlying origin of this failure to productively bind Rad4/yeast XPC to the site of the DNA lesion is the strong B[*a*]P aromatic ring system-DNA base stacking interactions that stabilize the G*:Del duplexes. Furthermore, the absence of the partner base[27] prevents the required flipping of that base to bind with specific amino acids in the protein. Similar structural and dynamic differences were delineated by MD simulations for the pair of UV-derived lesions, 6-4PP and CPD.[7] These are excellent and poor NER substrates in cell extracts, respectively, when positioned opposite their normal Watson-Crick partner bases. It is notable that these NER activities are also correlated with high (6-4PP) and low (CPD) Rad4/yeast XPC binding affinities.[7] The resistance of CPD lesions to NER in human cell-free assays is correlated with an XPC dissociation constant that is somewhat greater than the unmodified DNA value. MD simulations showed that 6-4 PP exhibited initial binding characteristics that can lead to productive binding, similar to the *cis*-B[*a*]P-G*:C full duplex, while the CPD lesion did not.[7]

**Figure 6.**
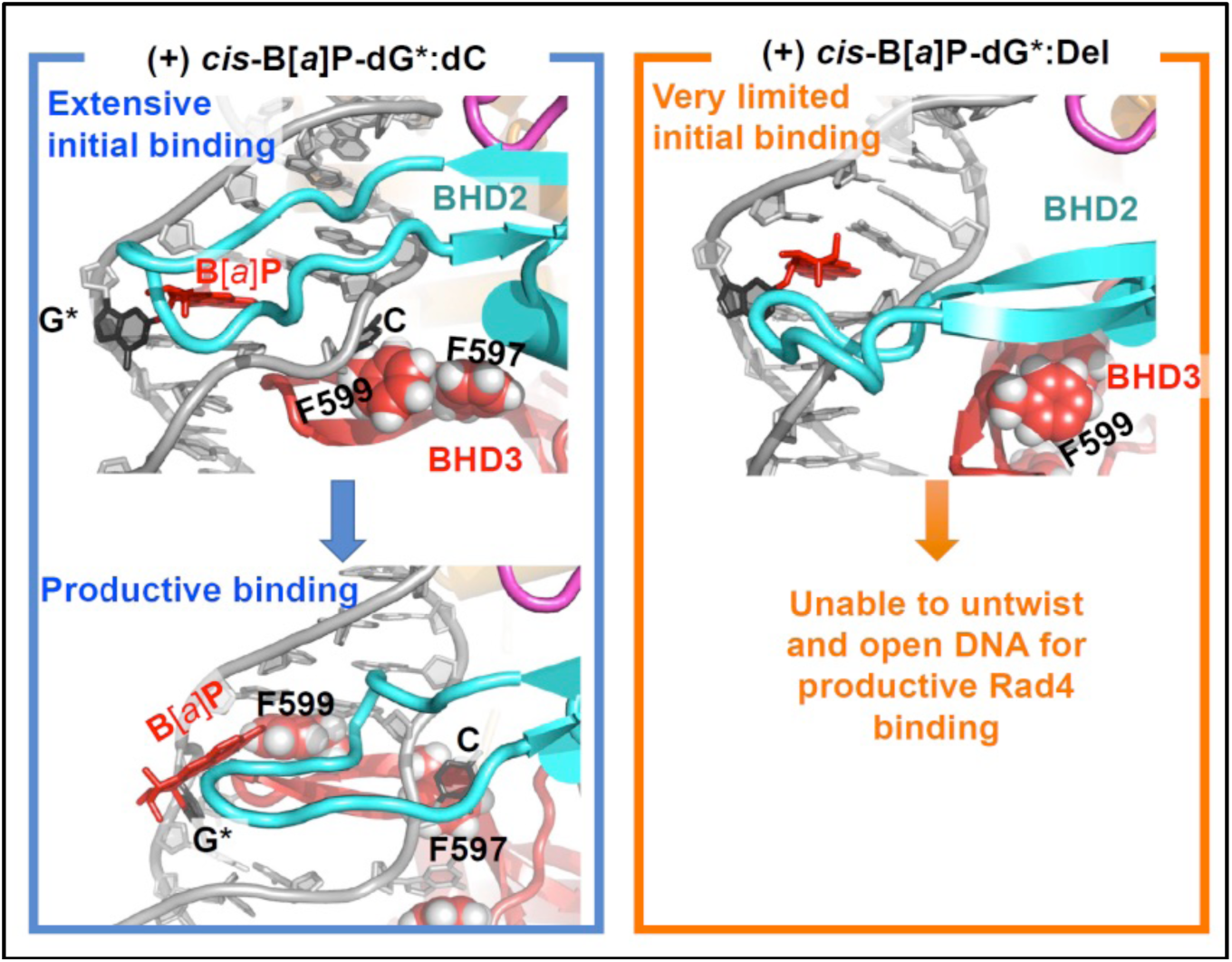
Structural origins of the productive binding of Rad4/yeast XPC to (+) *cis*-B[*a*]P-dG*:C and avoidance of binding to the (+) *cis*-B[*a*]P-G*:Del duplex, delineated by MD studies. The initial binding state structure for the (+) *cis*-B[*a*]P-G*:C case shows extensive initial binding by BHD2 which leads to lesion extrusion and BHD3 insertion to achieve productive binding. In the initial binding state of the (+) *cis*-B[*a*]P-G*:Del duplex, binding by BHD2 is very limited and the DNA cannot untwist and open to permit BHD3 insertion to achieve productive binding. The view is into the minor groove, looking at the BHD2 ß-hairpin. BHD3 is in the back, inserted from the major groove side in the productive binding state. The key phenylalanine (F599) that is inserted is denoted. BHD2 is in cyan, and BHD3 is in red.

Since binding affinities alone do not provide any information on the structures of the complexes, it is possible that the XPC-DNA lesion bound state(s) could be either productive and lead to excision, or non-productive and not excisable. A recent crystal structure[48] and previous experimental work[12] showed that there are bound states that do not form the productive open complex. Hence discrepancies between binding affinity and NER efficiency may result in part from non-productive binding and/or due to varying helicase-dependent verification efficiencies associated with the downstream NER factor TFIIH.

## 5. Conclusions

The low XPC binding affinities to the B[*a*]P-modified [G*:Del] duplexes offer new insights into the impact of base sequence context on NER-resistance that is consistent with an XPC lesion avoidance mechanism. This finding explains why the G*:Del lesions are resistant to NER.[24] Such a correlation is not observed in the case of the stereoisomeric B[*a*]P-dG adducts in full G*:C duplexes since the NER dual incision efficiency of the *cis*-B[*a*]P-dG adduct is ∼ 5 times greater in cell-free assays than the (+)-*trans*-B[*a*]P-G adduct [3] and 3.7±1 times greater when transfected into human cells.[49] The bulky B[*a*]P-dG adducts in deletion sequence contexts constitute a clear example of a correlation between full NER-resistance[24, 38] and weak XPC binding (*K*_*D*_(G*:Del)/*K*_*D*_(G:C) ∼ 3.2 ± 0.6 (*trans*) and ∼ 2.4 ± 0.3 (*trans*), Table 1. These results suggest that a partially non-productive XPC binding to the *trans*-adduct, or the verification step, or other still unknown effects, contribute to the overall difference in NER efficiencies of excision of the *trans*- and *cis*-B[*a*]P-dG adducts in full G*:C duplexes.

## Acknowledgement

This work was supported by NIEHS grant 1R01ES024050 (N.E.G.) and in part by 1R01ES027059 (V.S.), and R01-ES025987 to S.B. This work used the Extreme Science and Engineering Discovery Environment (XSEDE), which is supported by National Science Foundation (NSF) Grant MCB-060037 to S.B., and the high performance computing resources of New York University (NYU-ITS), as well as by National Science Foundation (NSF) grant MCB-1412692 (to J.-H.M) and National Institutes of Health grant (R21-ES028384 to J.-H.M).

