## Supporting Information for "The DNA Damage-Sensing NER Repair Factor XPC-RAD23B Does Not Recognize Bulky DNA Lesions with a Missing Nucleotide Opposite the Lesion"

### Supporting Figures

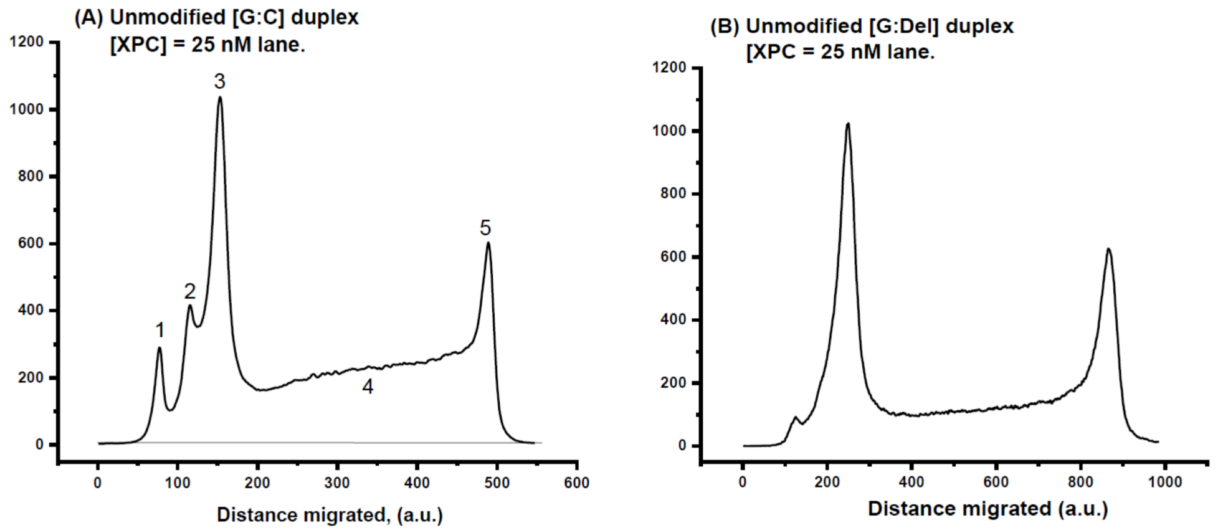


**Figure S1. Densitometry tracings of selected [XPC] = 25 nM lanes from the autoradiographs shown in Fig. 2 in the main text.**

(A) Unmodified 50-mer [G:C] duplexes: (1) well. (2) $(XPC)_2$-[G:C] DNA complexes. (3) (XPC)-[G:C], single XPC-DNA complexes.(XPC)-DNA. (4) Signals due to dissociated (XPC)-[G:C] and $(XPC)_2$-[G:C] complexes. (5) Free DNA.

(B) XPC binding to Unmodified (XPC) = [G*:Del] duplexes. No directly visible $(XPC)_2$-[G*:Del] in contrast to pane; (A), and the fraction of unstable complexes is also lower than in panel (A).

### Supporting Methods

#### Curve fitting calculations.

The exact analytical solution of expression characterizing competitive binding of two different DNA molecules (unmodified competitor G:C duplexes and BPDE modified DNA duplex (G*:C) in our case) (Eq. **S1-S5**), was derived by Wang.

According to the laws of mass conservation:

$$[G{:}C]_0 = [G{:}C] + [P \cdot (GC)] \quad \textbf{(S1)}$$

$$[G^*{:}C]_0 = [G^*{:}C] + [P \cdot (G^*C)] \quad \textbf{(S2)}$$

$$[P]_0 = [P] + [P \cdot (G{:}C)] + [P \cdot (G^*C)] \quad \textbf{(S3)}$$

where **[G:C], [G*:C]** and **[P]** are the concentrations of free unmodified, free modified DNA duplexes, and free protein concentrations, respectively.

Substitution and rearrangement of (**4,5,** main paper) and (**S1, S2**) into **(S3)** gives the following cubic equation in [P]:

$$[P]^3 + a\,[P]^2 + b\,[P] + c = 0 \quad \textbf{(S4)}$$

where

$$a = [G{:}C]_0 + [G^*{:}C]_0 + K_D(G{:}C) + K_D(G^*{:}C) - [P]_0 \quad \textbf{(S5)}$$
$$b = K_D(G^*{:}C\ ([G{:}C]_0 - [P]_0) + K_D(G{:}C\ )([G^*{:}C]_0 - [P]_0) + K_D(G{:}C)\ K_D(G^*{:}C) \quad \textbf{(S6)}$$
$$c = -\ K_D(G{:}C)\ K_D G^*{:}C\ [P]_0 \quad \textbf{(S7)}$$

By using proper substitution for eliminating the quadratic term, and general techniques of solving cubic equations, the only physically relevant root of equation **(S4)** can be written as

$$P = 2/3\ \sqrt{a^2 - 3b}\ \cos\theta - a/3 \quad \textbf{(S8)}$$

where

$$\theta = \frac{\cos^{-1}(-2a^3 + 9ab - 27c)}{2(a^2 - 3b)^{3/2}} \quad \text{(S9)}$$

and the fraction of the B[*a*]P-modified oligonucleotide-XPC complexes is:

$$y = P/(K_D + (G^*{:}C) + P) \quad \textbf{(S10)}$$

The XPC binding constants for B[*a*]P-G*:C duplexes were experimentally determined utilizing a 25-fold greater concentration of unmodified G:C duplexes (see Fig.3 of the main paper).

Analytical expressions for the binding of dimeric $(XPC)_2$-G*:C to DNA were analyzed by the (generic) mathematical methods described in [2,3]. The analytical expression for such equilibria is derived via the solution of cubic equations for the free protein concentration, that is similar to equations (**S4-S9**) for the competitive XPC-G*:C binding case because the same two protein molecules are involved in both cases, but the values of the coefficients (**S5-S7**) of the cubic equation (**S4**) are different.

Nonlinear curve fitting of the XPC binding to unmodified oligonucleotides gives the values of $K_D$(G:C), which in turn, were used for the nonlinear fitting of the competition binding data using equations (**S5-S10**), and the determination of $K_D$(G*:C) that provides the best calculated fit to the experimental data points.

------------------------------------------------------------
